# A novel enhancer RNA, Hmrhl, positively regulates its host gene, *phkb,*in Chronic Myelogenous Leukemia

**DOI:** 10.1101/378984

**Authors:** Roshan Fatima, Subhendu Roy Choudhury, T. R. Divya, Utsa Bhaduri, M. R. S. Rao

## Abstract

Noncoding RNAs are increasingly being accredited with key roles in gene regulation during development and disease. Here we report the discovery and characterization of a novel long noncoding RNA, Hmrhl, which shares synteny and partial sequence similarity with the mouse lncRNA, Mrhl. The human homolog, Hmrhl, transcribed from intron 14 of *phkb* gene, is 5.5kb in size, expressed in all tissues examined and has acquired additional repeat elements. Analysis of Hmrhl locus using ENCODE database revealed that it is associated with hallmarks of enhancers like the open chromatin configuration, binding of transcription factors, enhancer specific histone signature etc. in the K562 Chronic Myelogenous Leukemia (CML) cells. We compared the expression of Hmrhl in the normal lymphoblast cell line, GM12878, with that of K562 cells and lymphoma samples and show that it is highly upregulated in leukemia as well as several cases of lymphoma. We validated the enhancer properties of Hmrhl locus in K562 cells with the help of Luciferase assay. Moreover, siRNA mediated down-regulation of Hmrhl in K562 cells leads to a concomitant down regulation of its parent gene, *phkb*, showing that Hmrhl functions as an enhancer RNA and positively regulates its host gene, *phkb,* in chronic myelogenous leukemia.

## Introduction

Genome projects in recent years have revealed the fact that mammalian genome is transcribed virtually in entirety, but only a fraction of it is translated^1^, thus generating a plethora of noncoding RNAs (both long and short noncoding RNAs), the functions of a vast majority of which are yet to be determined. There has been a recent spurt in the studies of long noncoding RNAs (lncRNAs), demonstrating diverse roles not only in gene regulation during development and disease but also in evolution and complexity of organisms^2–3^. Enhancers constitute one such category of regulatory noncoding sequences that enhance the expression of their coding counterparts irrespective of their position, orientation and distance^4–6^. Any variation in enhancer sequence could lead to altered gene expression and consequent disease conditions^7^. Besides, super enhancers have been identified which consist of clusters of enhancers that are mainly associated with genes that define cell identity^8–10^. Recently, it has been observed that enhancers are actively transcribed giving rise to enhancer RNAs (eRNAs), making them the latest addition to the ever expanding list of regulatory noncoding RNAs. The eRNAs play a critical role in enhancer-promoter looping to bring about the activation of neighbouring protein coding genes^11–14^. In fact the enigmatic association of the gene desert loci with cancer has now been attributed to the enhancer derived long noncoding RNAs transcribed from the gene desert loci. For example, the 8q24 gene desert locus is known to give rise to several tissue specific long noncoding RNAs in different human cancers. CCAT1-L (Colorectal Cancer Associated Transcript-1) is one such enhancer RNA transcribed from a super enhancer region of 8q24 gene desert and has been shown to mediate long range chromatin interactions of Myc oncogene with its enhancers^15^. Further, a set of lncRNAs named as CARLos (Cancer-Associated Region Long noncoding RNAs) is also known to be transcribed from the 8q24 gene desert region. One of them, CARLo5 is an oncogeneic lncRNA associated with increased susceptibility to cancer and has been shown to be involved in long range chromatin interactions between Myc enhancer region and the promoter region of CARLo-5^16^.

Our lab had previously identified an intronic long noncoding RNA, Mrhl (Meiotic recombination hot spot locus RNA) in mouse^17^. The Mrhl lncRNA is 2.4kb in size and expressed in multiple tissues in mouse and is shown to be nuclear restricted^18^. Diverse roles have been attributed to Mrhl in cellular processes and signalling pathways. For example, it has been reported that Mrhl negatively regulates Wnt signalling in mouse spermatogonial cells through its interaction with p68 RNA helicase^19^. Conversely, the regulation of Mrhl by Wnt signalling has also been reported^20^. Moreover, genome wide chromatin occupancy of Mrhl and its role in gene regulation has been explored^21^. Very recently, the role of Mrhl in regulation of Sox8 expression and meiotic commitment of spermatogonial cells has been described^22^. In the current communication, we report the identification and characterization of its human homolog, Hmrhl, and show that Hmrhl functions as an enhancer RNA for its host gene, *phkb,* in K562 Chronic Myelogenous Leukemia cells.

## Results

### Syntenic Conservation of Mrhl in humans within the *phkb* gene

The Mrhl long noncoding RNA is transcribed from the intron 15 of the *phkb* gene located on chromosome 8 in mouse (Fig1.i). Noncoding RNAs are poorly conserved across species and it had earlier been reported from our lab that the Mrhl lncRNA is not conserved in human^18^. However, a low stringency blast of Mrhl sequence revealed that it is partially conserved in human within the same host gene, *phkb*, thus sharing synteny (Fig1.ii). The human homolog is transcribed from the intron 14 of the *phkb* gene (with respect to transcript # 201, the longest transcript of *phkb*, ENSEMBLE genome browser; Fig1.iii). The sequence from the middle region of Mrhl to the 3’end is conserved in human as two segments, with 71% identity over a stretch of 152bp and 65% identity over a stretch of 1223bp (Fig.1.iv-v). Therefore, the mouse Mrhl long noncoding RNA shares both synteny and partial sequence similarity with its human homolog, which we have named as human Mrhl or Hmrhl.

**Fig.1.**
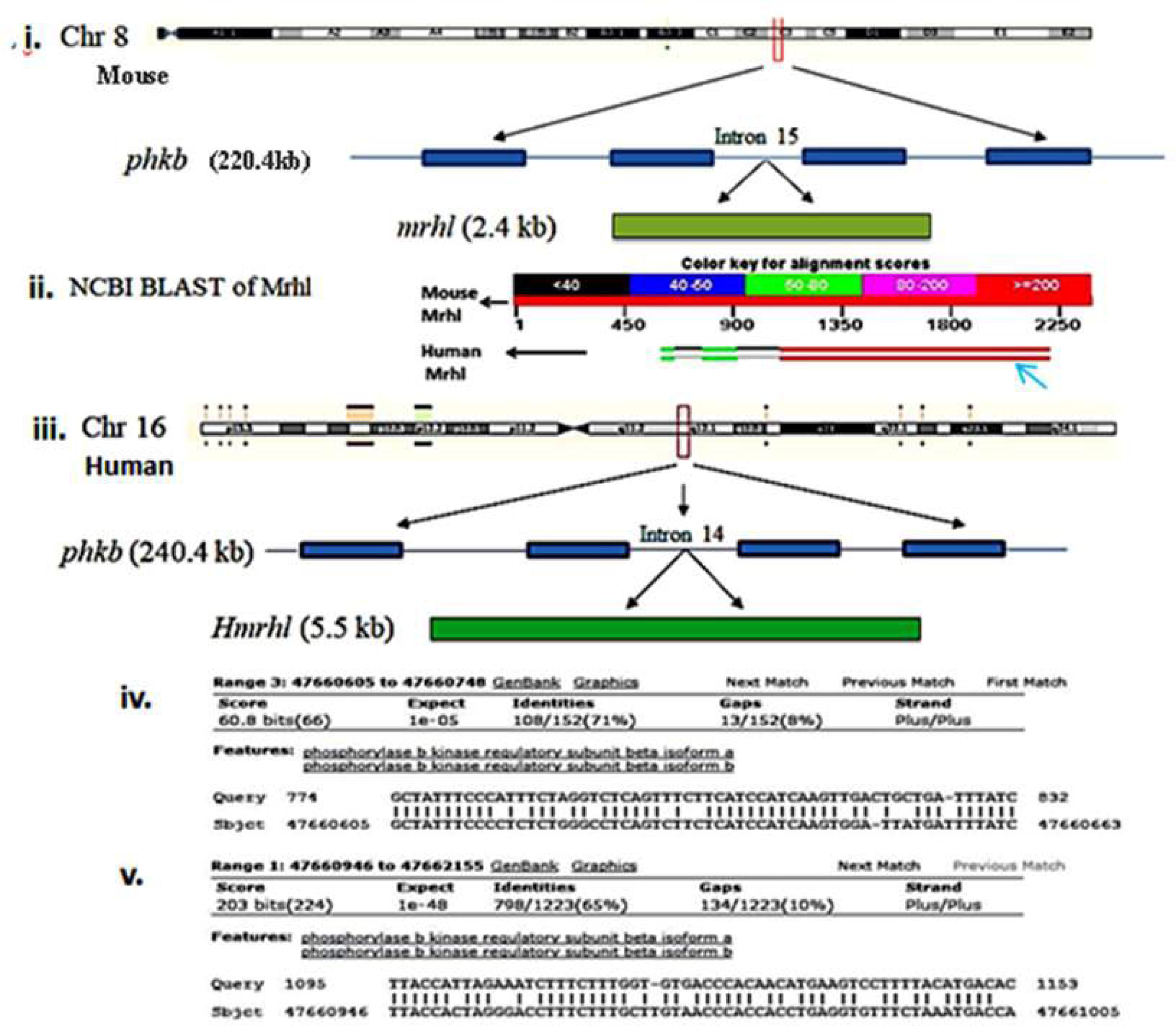
Blast Analysis of mouse Mrhl lncRNA id en tif i ed th e h u man h omol og. **i.** Schematic representation of mouse chromosome 8 and the C3 cytogenetic region which harbours the *phkb* gene. Mrhl is transcribed from the intron 15 of mouse *phkb* gene (shown as a pale green bar). **ii.** Blast analysis of mouse Mrhl identified the homologous sequence in the human *phkb* gene. Sequence from the middle region to the 3’end of Mrhl showed high homology with its human counterpart, as indicated by the red bars (Blue arrow). **iii.** Schematic representation of Human chromosome 16 and the 16q12.1 cytogenetic region where the *phkb* gene is located. Hmrhl (shown in deep green color) is transcribed from the intron 14 of the human *phkb* gene. **iv-v.** Sequences in the human genome sharing homology with Mrhl (GrCh37). Note the stretches of 71% (iv) and 65% identities (v) of the query sequence, Mrhl, against the human *phkb* gene (only part of the sequence shown here).

### Hmrhl is expressed in multiple human tissues

Once it became clear that the Mrhl sequence is conserved in human, we were curious to know whether the human homolog is expressed. For this, a cDNA panel of 16 different human tissues was obtained (Clontech, USA) and used to examine the expression of Hmrhl by quantitative Real Time PCR (qRT-PCR). Glycerldehyde-3-phosphate dehydrogenase (GAPDH) was used as an internal control and the data was plotted following the standard 2^-ΔΔCT^ method. Low level of Hmrhl expression (high cT values) was indeed observed in all of the human tissues examined (Fig.2a). Among all, the Hmrhl expression was highest in Spleen and Pancreas followed by Testis and other tissues (Fig.2a).

**Fig.2.**
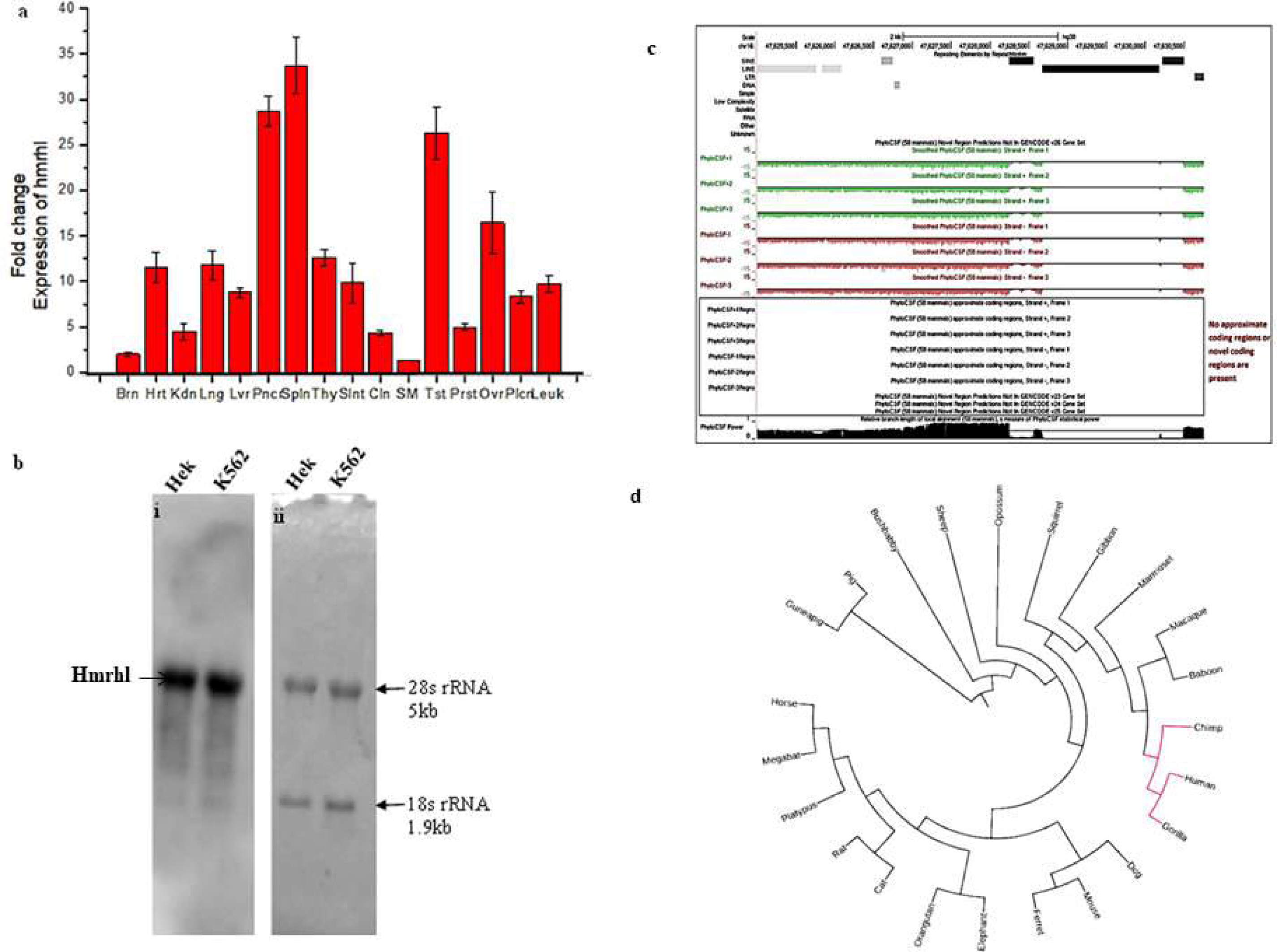
Expression and coding potential analysis of Hmrhl. **a.** Quantitative real time PCR analysis of Hmrhl expression showed that it is expressed in all human tissues (Brain, Heart, Kidney, lung, liver, pancreas, spleen, thymus, small intestine, colon, skeletal muscle, testes, prostate, ovary, placenta, leukocyte, from left to right) examined. Lowest expression was found in skeletal muscle (SM) which was taken as control, the level of which was considered as 1 and all others were plotted in comparison to it. Highest expression was seen in spleen (spln) followed by pancreas (Pnc), testis (Tst) and other tissues. **b.** Northern blot detection of Hmrhl. Total RNA from HEK 293T and K562 cell lines were separated on a 2% agarose gel and subsequently hybridized with DIG labelled Hmrhl specific riboprobe to detect the transcript (i). In parallel, methylene blue staining was used to determine the size of HMRHL, using 28S rRNA (5kb) and 18s rRNA (1.9 kb) as reference (ii). Note that the size of Hmrhl is similar to that of 28s rRNA, revealing that Hmrhl is about 5kb in size. **c.** Protein-coding potential as determined by Broad Institute’s PhyloCSF data and visualized in UCSC Genome Browser, showing that Hmrhl has no coding potential. **d.** Circular phylogenetic tree built in iTOL (Interactive Tree of Life).

### Hmrhl is 5523bp in size, much larger than its mouse homolog

For a detailed characterization of Hmrhl lncRNA, we used the most commonly available commercial cell line, human embryonic kidney cells (HEK293T) as well as the K562 CML cells. Hmrhl sequence was cloned in pGEM3Zf(+) vector to generate a template for a northern probe. The Hmrhl specific probe detected a single band in the northern blot indicating the presence of the Hmrhl transcript (Fig.2b), the size of which was much larger (~5kb) as compared to Mrhl (2.4kb). In order to determine the exact size of Hmrhl lncRNA, Primer Walking method was further employed to reach the ends of Hmrhl cDNA from the middle conserved region. Commercially available panel of human cDNA of different tissues (Clontech, USA), as well as the HEK293T cDNA was used in combination with different overlapping sets of primers (Fig.S1). This experiment revealed the size of Hmrhl to be 5523bp, corresponding to chr16: 47658923-47664446 on GRCh37/ hg19, the genome build available at the time of mapping and chr16: 47625012-47630534 on GRCh38/ hg38, the current genome build. The sequence for Hmrhl is submitted to the NCBI (Genbank ID MF784605, FigS2).

We next investigated the coding potential of Hmrhl with NCBI ORF Finder which showed the presence of 1510bp long ORF (Fig.S3), though its translation is highly unlikely as discussed in supplementary data. To further examine the plausible coding potential of Hmrhl, the online tools CPAT analysis (Fig. S4) and PhyloCSF were used (see methods section for details) showing and strongly advocating that no approximate coding regions or novel coding regions are present (Fig.2c).

Syntenic conservation of Hmrhl allowed us to further explore its sequence conservation across species through evolution. Phylogenetic tree for Hmrhl was generated using Circular phylogenetic tree built in iTOL (Interactive Tree of Life), as shown in Fig.2d. Among the twenty-three-selected species, the Hmrhl sequence seems to have its closest similarity with the conserved region present in the Gorilla followed by Chimp, although the single exonic Hmrhl carries a definite region that appears to be highly conserved among primates.

### Analysis of Hmrhl locus using ENCODE portal

Data from the ENCODE Project Consortium^1^ (visualized through the UCSC genome browser) was analysed exhaustively with respect to the Hmrhl cytogenetic region and the parent *phkb* gene to gain an insight into the structural and functional properties of Hmrhl. Remarkably, the ENCODE database revealed several interesting facts about Hmrhl genomic interval as discussed below.

### Hmrhl has acquired several repeat elements during evolution

A comparison of the genomic sequences in and around Hmrhl region from lower to higher vertebrates shows that the sequence appears to have been acquired initially in lower mammals like platypus during evolution and then evolved rapidly by insertion of repeat elements, especially in primates. Hmrhl has acquired seven different repeat elements in humans, L2b, L2c, MIR, Charlie 15a, AluY, L1PA3 and AluSx (Repeat Masker), which flank the unique central region (Fig3). While the central region is highly conserved from lower to higher mammals, the repeat elements present in Hmrhl are absent in lower organisms, including rodents. The mouse homolog has only one repeat element, RLT, which is not present in Hmrhl (data not shown).

**Fig.3.**
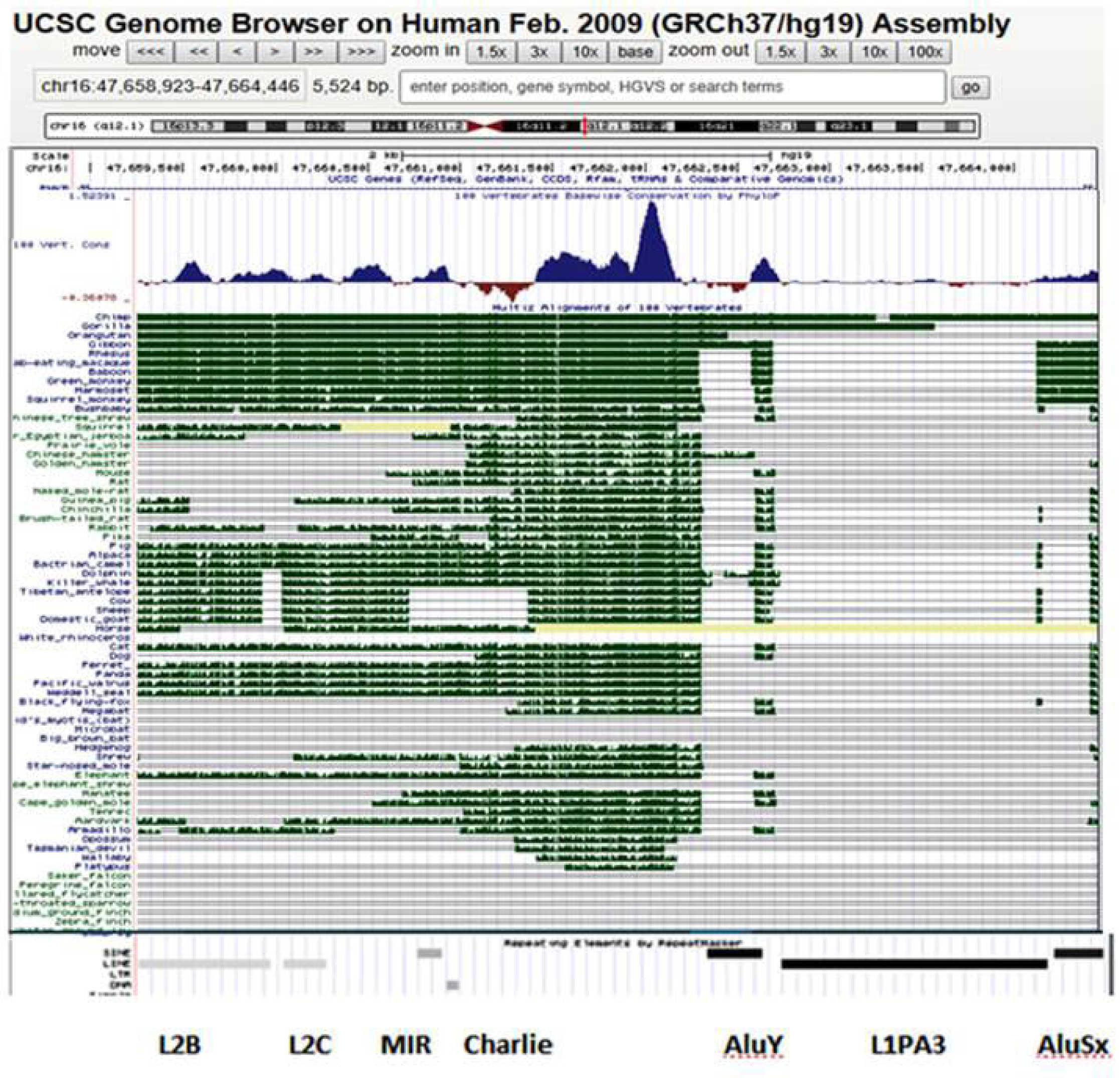
Hmrhl is conserved in mammals and has gained several repeat elements in the primates. Figure shows the conservation of Hmrhl across various organisms and the repeat elements present in this genomic interval (shown below the conservation tracks) in human, as visualized through the UCSC genome browser. Note the unique sequence in the middle of Hmrhl which is highly conserved across various organisms and is flanked by the repeat elements L2B, L2C, MIR, Charlie, AluY, L1PA3, AluSx which are present only in higher primates (see text for details).

### Hmrhl locus exhibits hallmarks of enhancer in K562 CML cells

Encode database represents a true encyclopaedia of DNA elements and offers valuable information regarding various properties of the chromatin like DNAse hypersensistive sites, transcription factor binding sites and transcriptional status, histone signature etc., across different cell types; through diverse organisms.

Interestingly, ENCODE database shed light on the fact that Hmrhl locus exhibits hallmarks of enhancers in the Chronic Myelogenous Leukemia cell line, K562. Hmrhl locus is associated with open chromatin configuration and the presence of DNAse hypersensitive sites in the K562 cells but not so in the normal lymphoblast cell line, GM12878 (Fig.4a). Moreover, Hmrhl locus exhibits the binding of EP300, histone signatures H3K27Ac and H3K4Me1 that mark the enhancers as well as the H3K4Me3 signal, which marks the promoters, in K562 cells (Fig.4a). Particularly, the H3K27Ac enhancer mark is in the form of two distinct peaks at the 5’ end of Hmrhl (Fig.4a). We validated these enhancer marks at Hmrhl locus using ChIP-Seq (Fig4 b-c). Furthermore, Chip-seq data from ENCODE shows the binding of RNA polymerase II and an array of transcription factors, namely, GATA1, PML, CCNT2, NR2F2, TRIM28, STAT5A, ATF1, cMyc, GATA2, IRF1, JUND, EGR1, RCORI, BHLHE40, YY1, TAL1, REST, TEAD4, MAFF, MAX and Egr1 at the 5’ end of Hmrhl (genomic co-ordinates ~chr16:47,658,800-47,659,606) in the K562 cell line (Fig.5a). Notably, most of these transcription factors have been linked to cancer in previous studies^23–25^. It is well known that transcription factors initially bind to enhancer elements and recruit RNA PolII and co-activators to target gene promoters through looping ^26–31^. EP300 is a histone acetyl transferase, a transcriptional coactivator and a chromatin re-modeller known to be enriched at active enhancers and to increase gene transcription by relaxing the chromatin and recruiting the transcriptional machinery^32–33^. All these observations strongly suggested that Hmrhl locus is transcriptionally highly active in K562 leukemic cells and that it possesses regulatory properties.

**Fig4.**
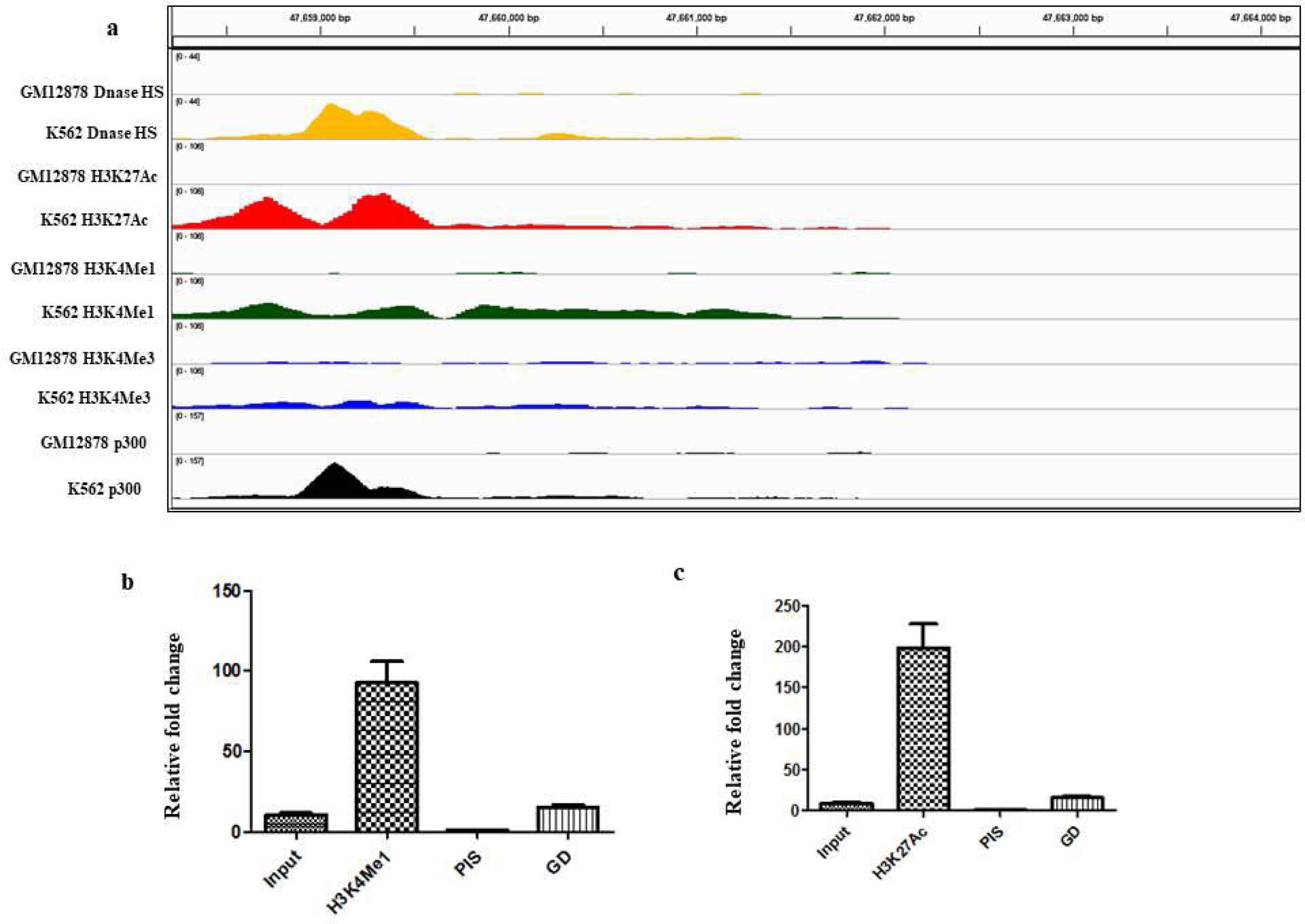
Hmrhl locus exhibits hallmarks of enhancer. **a.** ENCODE data visualized through Integrated Genome Viewer (IGV) for DNase hypersensitive sites, p300 binding, enhancer specific histone marks, H3K27Ac and H3K4Me1 and the promoter specific histone mark, H3K4Me3 at the 5’ end of Hmrhl, only in K562 but not in GM12878 cells. Note the two prominent peaks (red) for the enhancer mark H3K27Ac in K562. **b-c.** Chromatin immunoprecipitation with Ab4729 (anti-H3K27Ac antibody) and Ab8895 (anti-H4K4Me1 antibody) in K562 cells. Note the enrichment of both the enhancer marks at the 5’ end of Hmrhl in the IP fraction as compared to input/ PIS/ gene desert region (GD), that serves as a negative control.

One of the most significant disclosures of the Encode portal with respect to the Hmrhl is the fact that the Hmrhl locus loops and interacts with the promoter of its parent gene, *phkb,* in K562 leukemia cells, thus acting as an enhancer for *phkb* gene. This fact was revealed through the ChiaPET (Chromatin interaction analysis with Paired End Tag sequencing, a combination of Chipseq and 5C, Fig.5b-d) data of ENCODE project.

With this background information regarding Hmrhl locus from the Encode database as described above, we set out to elucidate its functional properties. We examined the expression profile of Hmrhl across various human cancers using a cancer specific cDNA panel (Origene, USA) by real time qPCR. Actin was used as an internal reference control. We observed different degrees of variation in the level of Hmrhl expression between normal and cancer samples under various cancers conditions. Significantly, it was observed that Hmrhl was highly upregulated in a number of lymphoma samples suggesting an active role for this RNA in solid tumors of blood cancer (Fig.6a). In addition, we compared the expression of Hmrhl and PHKB in the GM12878 and K562 cells by real time qPCR and found that both are over expressed in K562 leukemia cells as compared to normal lymphoblast (Fig.6 b-c).

**Fig.5.**
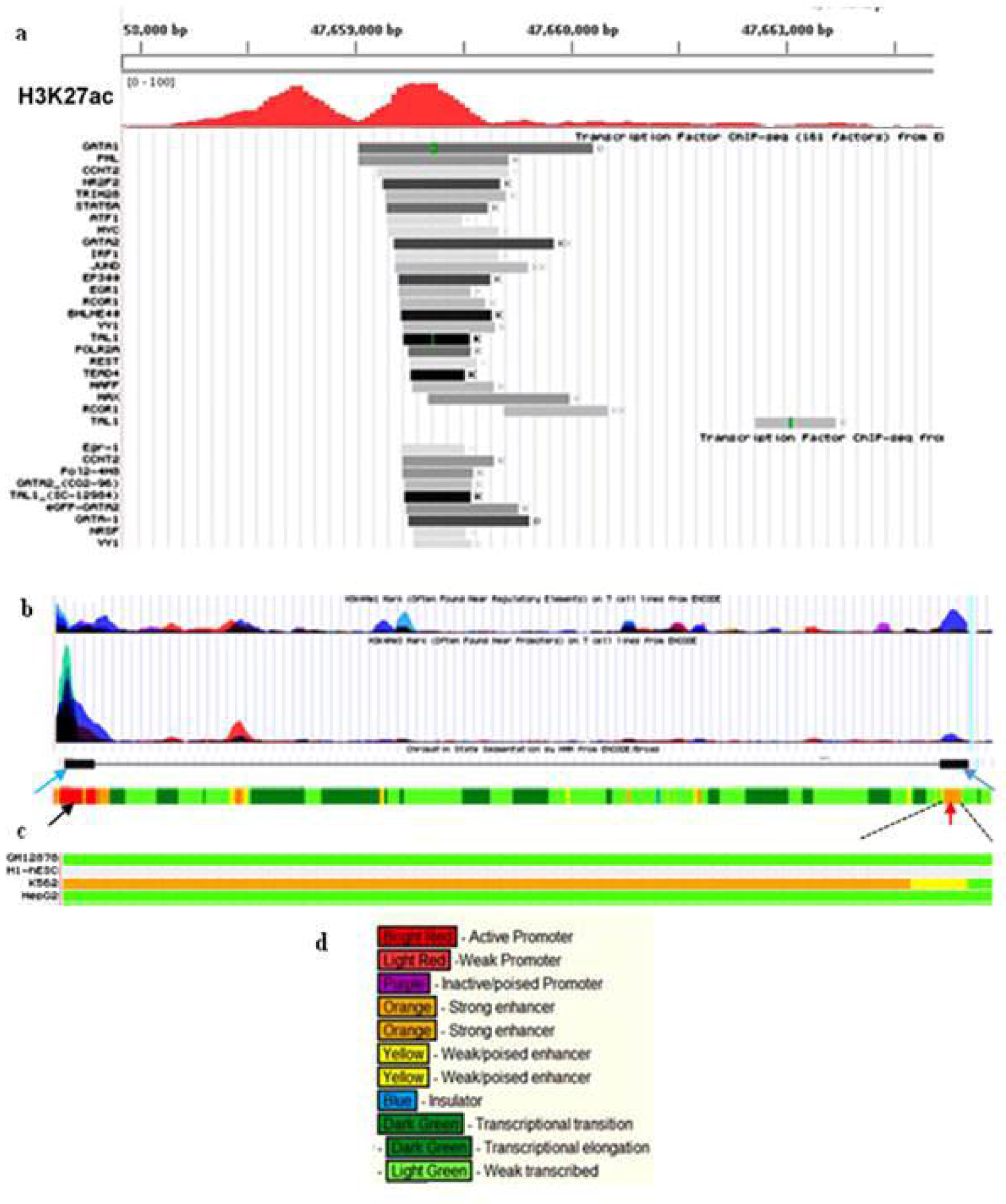
Hmrhl locus exhibits hallmarks of enhancer contd. **a.** Encode data shows the binding of various transcription and PolII at the 5’ end of Hmrhl. We have retained the H3K27Ac peaks in this figure also for a reference. **b.** Schematic for chromatin interaction analysis (ChiaPET data) for Hmrhl. The large purple-black peak representing histone marks on the extreme left denotes the promoter of *phkb* gene while the small purple peak at the far right represents the 5’end of Hmrhl. ChiaPET data shows the interaction of Hmrhl locus with *phkb* promoter, as represented by two black boxes (blue arrows) connected by a black line in **b.** The Hmrhl locus is expanded below in **c**, showing that this locus has enhancer properties only in K562 cell line (orange-yellow color), but not in other cell lines like GM12878, HepG2 or hESC. Genomic segments are colour coded by ENCODE as denoted in **d**, with red colour signifying active promoter (*phkb* promoter at far left, black arrow in **b**) while orange colour represents active enhancer at Hmrhl locus at far right (red arrow in **b**).

**Fig.6.**
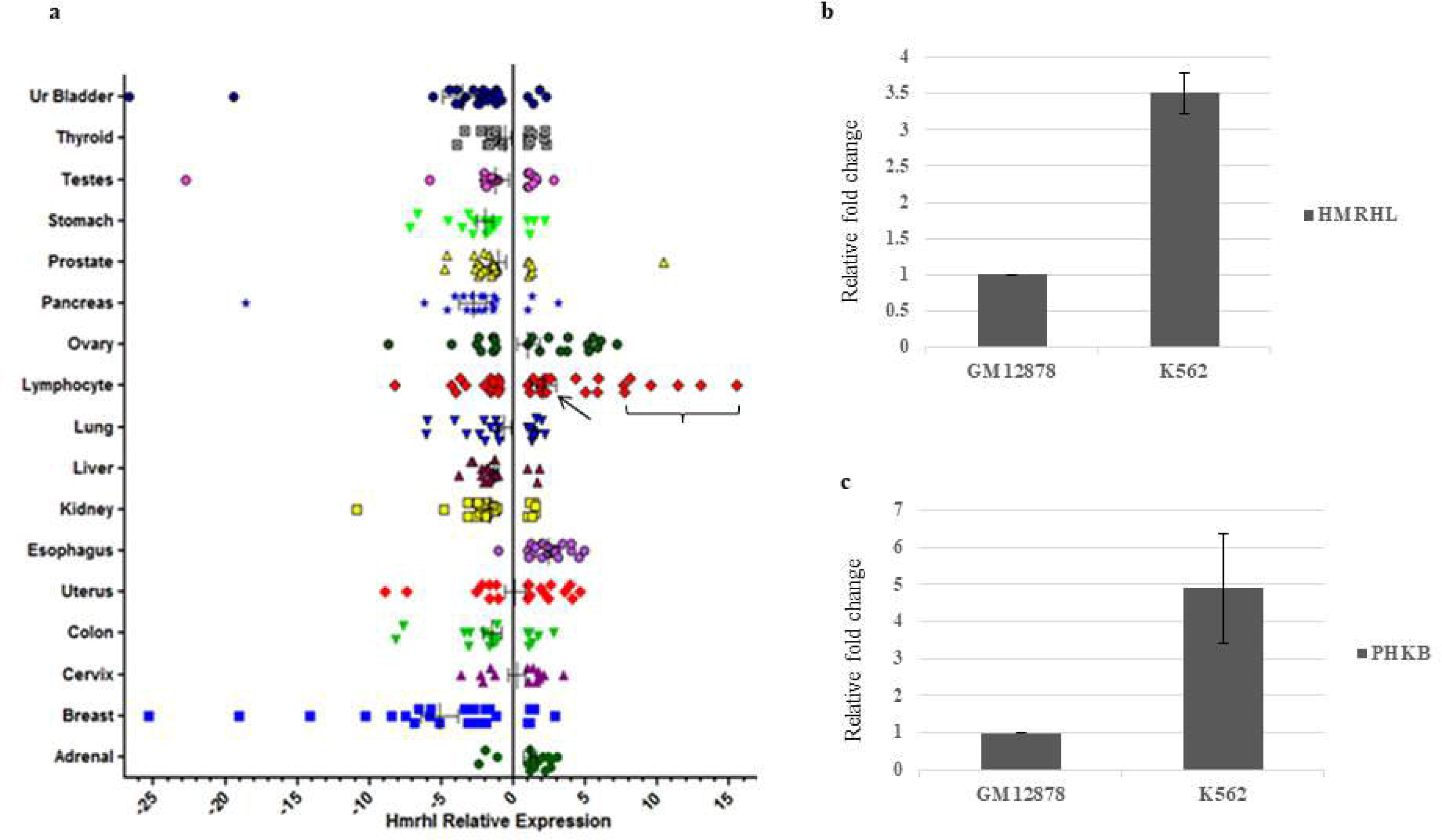
Hmrhl is differentially expressed in various cancers. **a.** Expression of Hmrhl in various normal and cancer samples as observed by qPCR. Note that Hmrhl is highly upregulated in several lymphoma samples (bracket) in comparison to normal range (arrow). In fact, of all cancers, the highest levels of Hmrhl are seen in some of the lymphoma samples. **b-c.** qPCR analysis of Hmrhl and PHKB expression showing that both are over expressed in K562 leukemia conditions as compared to GM12878 normal lymphocytes.

### Luciferase assay validated the enhancer properties of Hmrhl locus

In order to further ascertain the enhancer properties of Hmrhl genomic locus, we obtained the promoter and enhancer specific vectors pGL4.10 and pGL4.23, respectively (Promega, USA) for a luciferase assay. The pGL4.10 promoter vector does not possess any promoter, and hence can be used to identify promoter properties of any given DNA sequence when cloned upstream of the luciferase gene while the pGL4.23 enhancer vector has a minimal promoter upstream of luciferase gene and hence can be used to examine if a sequence of DNA has enhancer properties.

Three clones each for the promoter and enhancer vectors were generated using sequences in the 5’ end of Hmrhl as inserts. Insert-1 had a sequence of −1000bp upstream of the 5’end of Hmrhl. Insert-2 included a sequence of +1 to +500bp downstream of the 5’ end of Hmrhl (the region where various transcription factors and PolII were shown to bind by the Encode project), along with −300bp upstream sequence (800bp insert). Insert-3 included −1000bp upstream sequence and +500bp downstream sequences (1500bp insert). These inserts were cloned into the promoter and enhancer vectors, transfected in K562 cells and luciferase assay was performed to examine if any of these DNA sequences had regulatory properties.

Insert-1 did not generate any luciferase signal in combination with either promoter or enhancer vector, thus revealing that the −1000bp genomic sequence alone, upstream of the Hmrhl start site, has neither promoter nor enhancer properties. Insert-2 produced moderate signal both with promoter and enhancer vectors, suggesting that transcription factors and PolII indeed bound to this region, as also revealed in ENCODE database. Interestingly, an intense signal of luciferase activity was obtained with insert-3 (1.5kb) in combination with enhancer vector but not with promoter vector, confirming the fact that the sequences at the 5’ end of Hmrhl indeed have strong enhancer properties (Fig.7a).

**Fig.7.**
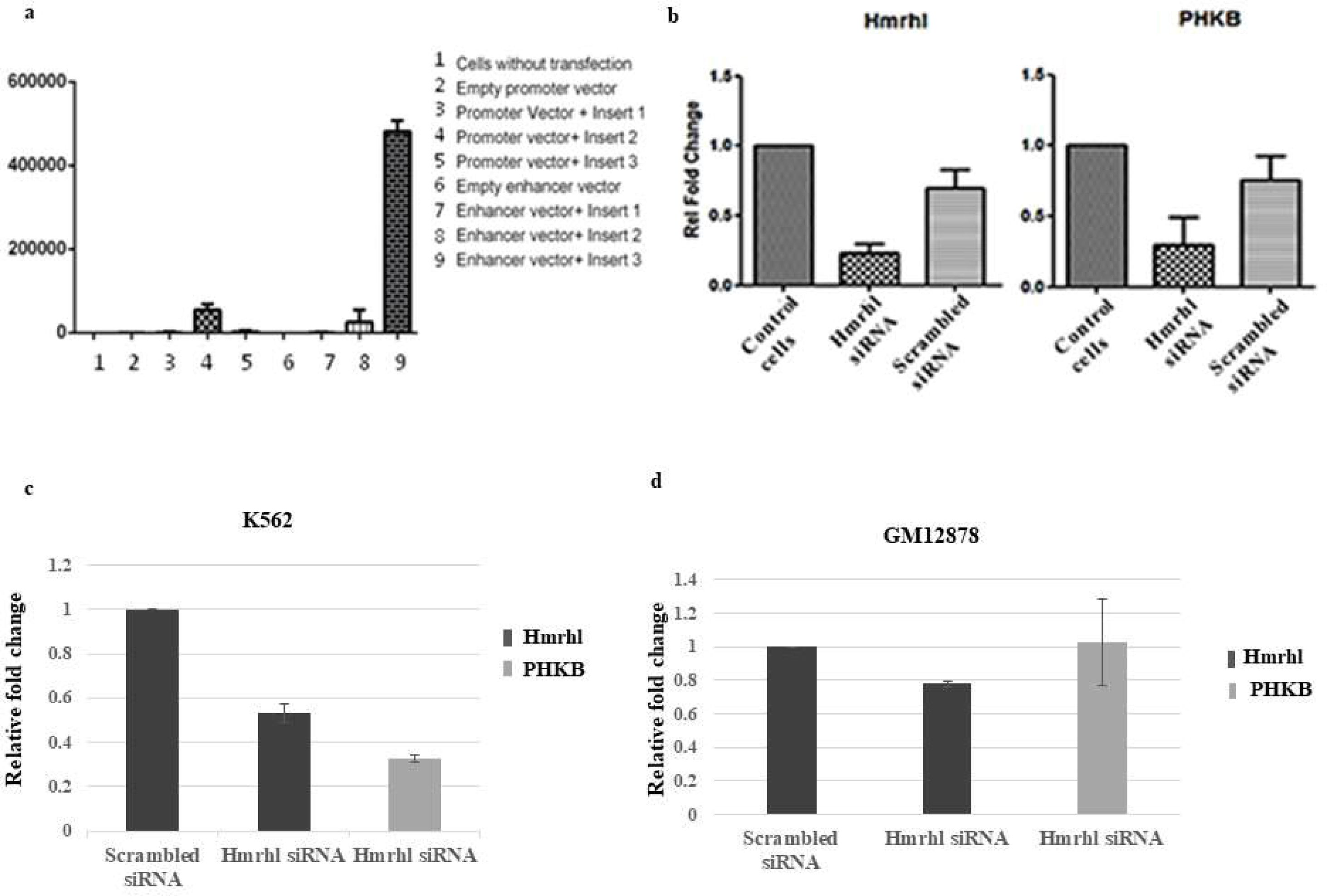
Hmrhl functions as enhancer RNA. **a.** Lucifaerase assay showing the intense signal of reporter activity in K562 cells with insert 3 cloned in enhancer vector. Note the low level of luciferase signal obtained with insert 2 both with promoter and enhancer vectors. **b.** siRNA mediated down-regulation of Hmrhl causes down-regulation of PHKB in K562 cells treated with Hmrhl specific siRNA pool as compared to control cells without transfection and cells treated with scrambled siRNA as negative control. **c-d.** Smart pool siRNA (Dharmacon) were used against the Hmrhl region to downregulate Hmrhl and subsequently expression level of PHKB gene were checked by qPCR in both K562 and GM12878 cell lines. Scrambled siRNA was used as a negative control.

### Hmrhl functions as an enhancer RNA for *phkb*

After confirming the enhancer properties of Hmrhl locus by luciferase assay, we went on to probe if only the genomic segment has the regulatory properties, or the transcript arising from this region also takes part in the positive regulation of its parent gene, *phkb*. Specific siRNAs against the Hmrhl lncRNA were obtained in order to silence it. A pool of four different siRNAs was used to down regulate Hmrhl in K562 cells and its effect on PHKB expression was examined through quantitative PCR. PHKB was indeed found to be down-regulated upon down-regulation of Hmrhl, confirming the fact that Hmrhl acts as an enhancer RNA and positively regulates its parent *phkb* gene in K562 cells (Fig.7b). This experiment clearly established the role of Hmrhl as an enhancer RNA involved in the regulation of its host protein coding gene. Moreover, using another set of siRNAs (targeting different region of Hmrhl), we observed the similar down regulation of PHKB upon knockdown of Hmrhl (Fig.7c). We further probed if the PHKB expression is affected upon downregulation of Hmrhl in the GM12878 normal lymphoblast cells as well and found that PHKB is not down regulated in GM12878 cells unlike in K562 leukemia cells (Fig.7 d).

## Discussion

Enhancers are mainly cis regulatory elements involved in the spatio-temporal regulation of gene expression. Enhancer derived transcripts were described initially way back in early 1990s when it was reported that the HS2 enhancer of K562 cells gives rise to long enhancer transcripts^34^. Yet there had been a long silence in the discoveries and understanding related to enhancers and their probable mechanism of action. In fact enhancers were perceived only as segments of DNA^4^ and the prospect of their transcription and a critical role for those transcripts in gene regulation was not envisaged. The discovery of enhancer RNAs came about rather recently, in 2010, with two papers describing the enhancer generated transcripts in mouse and human, respectively^11–12^. Since then, there has been a surge in studies related to enhancer RNAs which endorse the fact that enhancer function is mediated through enhancer transcripts, changing the way we appreciate mechanism of gene regulation^14, 35–40^.

In the present communication, we report the identification and characterization of the human homolog, Hmrhl, of the mouse long noncoding RNA, Mrhl and show that Hmrhl functions an enhancer RNA for its host gene. Both the Mrhl as well as Hmrhl lncRNAs are transcribed from the introns of the same protein coding gene, *phkb*. Remarkably, as many as 50% of human protein coding genes are known to act as hosts for noncoding RNA genes including micro RNAs, small nucleolar RNAs and lncRNAs^41^.

Recent reports have implicated a large number of long noncoding RNAs in the pathogenesis of various cancers^42^. Our current observations with experiments involving luciferase assay and siRNA mediated RNA interference, in the back drop of ENCODE data, clearly establish that Hmrhl functions an enhancer RNA and positively regulates its parental *phkb* gene.

Though siRNAs are known to have off target effects, the fact that two different pools of siRNAs generated the same effect, one of them being a smart pool from Dharmacon designed to target specific region of interest, the down-regulation of *phkb* can be held as a specific consequence rather than a non-specific off target effect. Further, given the fact that splicing is predominantly co-transcriptional^43–45^, it is highly unlikely that siRNAs directed against Hmrhl would affect the primary transcript through the intron leading to its down-regulation. In fact the presence of RNA PolII, transcription factors and active epigenetic marks at 5’ end of Hmrhl strongly suggest it to be an independent transcript, rather than being spliced out of/ arising from the PHKB primary transcript/ intron. This fact is substantiated by our northern hybridization and primer walking experiments. Northern hybridization with Hmrhl specific probe detected a single transcript of ~5 kb and no larger transcript of the size corresponding to the intron (30 kb) was recognized. Moreover, we reached ends of the Hmrhl transcript at 5523 bp itself during primer walking, confirming the fact that it is not a part of any larger, stable intron but rather an independent transcript. Therefore, we propose that the down-regulation of *phkb* upon knockdown of Hmrhl is related to regulation of *phkb* transcription by the Hmrhl enhancer RNA, rather than being mediated through the intron or the primary transcript.

The human *phkb* gene is located on the chromosome 16: 47,461,123-47,701,523 (240kb, GRCh38) at 16q12.1 cytogenetic region, on the forward strand. It is involved in the breakdown of glycogen to produce glucose. An up-regulation of *phkb* gene to produce more glucose is arguably beneficial for the cancer cells in order to sustain their increased metabolic rate and proliferation. It has indeed been reported recently that *phkb* gene promotes glycogen breakdown and aids cancer cell survival^46^. In fact PHKB shows medium to high levels of expression in several blood cancers as shown in cell line atlas data, with highest level of expression seen in K562 (supplementary Fig. S5).

The primary cause of chronic myelogenous leukemia is a translocation between chromosomes 9 and 22, which generates the shortened chromosome 22 known as Philadelphia chromosome^47^. This translocation creates a fusion oncogene, BCR-ABL, which codes for a constitutively active Tyrosine Kinase protein. This protein in turn activates a cascade of genes involved in cell cycle and inhibits those involved in DNA repair, thus leading to an uncontrolled growth of CML cells with an accumulation of secondary mutations^47^. Corroborating this, a very recent study by Zhou et al (2018)^48^ has revealed a catalogue of sequence, structural and copy number variations in K562 genome. They show that K562 genome is near triploid in nature, shows insertion of several novel LINE and Alu elements, multiple chromosomal translocations including the hallmark BCR/ABL translocation, and an array of indels, SNPs and other mutations. Further, genome wide association studies (GWAS) show that more than 90% of the disease associated SNPs fall in the noncoding portions of the genome and that many SNPs fall within or very close to enhancers^10, 49^. Cavalli et al (2016)^50^ sequenced a number of ENCODE cell lines and showed that highest number of the allele specific, disease associated SNPs were detected in K562. Previous studies have also revealed that cancer cells acquire de novo enhancers at driver genes. For example, Hnisz et al (2013)^10^ identified super enhancers associated with key oncogenes in 18 different cancers with the help of ChIP-seq for H3K27Ac mark. The gene desert region around the Myc oncogene acquires H3K27Ac enhancer mark in colorectal cancer, pancreatic cancer and T-cell leukemia, which is absent in the healthy individuals^10^. In view of all the above observations, we propose that Hmrhl locus has gained an enhancer/ super enhancer function in the K562 cells either due to change/s in the primary sequence or due to a change in the activity/ availability of factors like enzymes, transcription factors, looping factors etc., or due to a combination of many such events.

Our luciferase reporter assay revealed that insert 3 (1.5kb), which included both the −1KB upstream sequence as well as the +500bp downstream sequence of Hmrhl start site produced a very strong signal with the enhancer vector as compared to the other two inserts. In fact, of the two H3K27Ac peaks formed at the 5’ region of Hmrhl, one is downstream of the transcription start site of Hmrhl while the other is upstream of it. Interestingly, when we analysed −1kb Hmrhl upstream sequence, it was revealed that they possess sequences very similar to the core sequence of viral enhancers namely AAAACCAC and GTGGTTTGAA^51^ (Fig. S6). These viral enhancers are precisely conserved in the mouse as well as human immunoglobulin heavy chain gene enhancers^52–55^. Not only the B cells, even the T cells and haematopoietic cells in general, have specific viral enhancers which are located within introns^56–58^. Several viruses including the polyoma virus and the Molony murine leukemia virus (MoMLV) possess these enhancer elements and any mutations in the core sequence, ‘AAAACCAC’, have been shown to cause increased incidence of erythro leukemias^59–61^. A group of mammalian transcription factors involved in haematopoiesis called Core Binding Factor (CBF), binds to the core site of many retroviral enhancer elements and also to the enhancers of T-cell receptor genes^62–63^. One of the subunits of CBF, CBF-α, also known as AML1 (Acute Myeloid Leukemia 1), is known to be rearranged by chromosomal translocations in myeloproliferative diseases and mutations in core AML1 sites in murine leukemia viruses are known to affect their disease specificity and latency^59, 62, 64–66^. In all probability, the viral enhancer elements in the immediate upstream region of Hmrhl seem to play a critical role in conferring the enhancer properties to Hmrhl in K562 erythro-leukemia cells. In essence the viral enhancers could mediate the enhancer evolution in the human genome. Though our analysis of the Hmrhl locus and its 1kb upstream region did not reveal any variations in the sequence, it may be noted here that interrogating a complex cancer genome like that of K562 is not a simple task by any means. Since K562 genome is triploid in nature and bears multiple mutations, we advocate that a blend of different genetic/ epigenetic alterations could have rendered the enhancer properties to the Hmrhl locus.

The *phkb* gene appears to share the promoter with its upstream gene, ITFG1, but it remains to be seen if Hmrhl has a role in regulating ITFG1 gene as well, in leukemia.

Sequence conservation of noncoding regions is suggested to indicate the occurrence of enhancers^67^. In this context, it may be noted that both Mrhl and Hmrhl are transcribed from the introns of *phkb* gene and show significant sequence conservation from mouse to human as seen in current studies. The fact that even the mouse counterpart, Mrhl exhibits the enhancer marks, H3K4Me1 and HeK27Ac in the mouse erythro-leukemia cells (MEL cell line, mouse Encode database, Supplementary Fig.S7), strongly suggests a functional conservation of these two homologs, especially under leukemic conditions. Intronic enhancers/ eRNAs have been reported in case of several other genes as well ^39,68-69^. The eRNAs are known to be unstable and low in abundance^38–39^, possibly due to which they were not easily detected in earlier days, in the absence of high throughput techniques like RNA sequencing.

With respect to the mouse Mrhl function, our earlier studies in the mouse spermatogonial cell line, GC1-SPG had revealed that Mrhl down-regulation brings about the activation of Wnt signalling but does not affect its parent gene, *phkb*^19^. It was further reported that Mrhl gets down-regulated in response to induced activation of Wnt signaling^20^. It may be noted here that a human spermatogonial cell line is not available wherein the functions of Mrhl and Hmrhl could be aptly compared, with respect to the Wnt signalling regulation function. As far as the K562 cell line is concerned, Hmrhl is well expressed in these cancer cells despite the active Wnt signalling, rather than being downregulated, unlike what has been reported for its mouse counterpart in the GC1 cells^20^. We did not observe a translocation of β- catenin to the nucleus under conditions of Hmrhl downregulation in Hek293T cells either (Fig.S8). Even in case of mouse embryonic stem cells also, Wnt signalling regulation by Mrhl has not been observed (Pal et al, unpublished). These studies show that regulation of Wnt signalling by the Mrhl RNA is not a global phenomenon but it could be exclusive to the mouse spermatogonial cells. It may perform a completely different task in another cell type or in another cellular context, which appears to be the case with regard to the erythro leukemia cells.

Enhancers are regulatory elements that ensure tissue and developmental stage specific expression of genes, since all genes exist in all tissues, throughout development, but express only when/ where the enhancer is active. As mentioned earlier, genome wide association studies have revealed that enhancers are the prime targets for genetic and epigenetic changes that support cancer initiation and tumor progression. Mutations in enhancers or gain of super enhancers have been reported to favour cancer development in a number of cases^8,70-72^. A recent report by Corces Zimmerman et al (2014)^73^ describes super enhancers specifically in a group of AML patients. The targets of these super enhancers involve not only the key driver genes of AML but also genes encoding protein kinases and chromatin regulators, providing insights into the significance of super enhancers in the context of cancer.

Understanding related to enhancers and their mechanism of action is expected to advance diagnosis and therapeutic strategies for cancer and other diseases linked to altered enhancer function. Enhancer RNAs can serve as valuable biomarkers for various diseases and the expression of the causal genes can be manipulated through RNA interference mediated gene silencing for a promising remedy.

## Materials and Methods

### Cell lines and reagents

K562 (Chronic Myelogenous Leukemia/ Erythro Leukemia) cell line was obtained from NCCS Pune (India) and cultured in RPMI medium (Gibco), Hek293T cells were obtained from the American Type Culture Collection (ATCC, CRL-1573) and were cultured in Dulbecco’s modified Eagle’s medium (Sigma); both were supplemented with 10% fetal bovine serum (Invitrogen); and 100 units/ml penicillin-streptomycin solution (Sigma) at 37°C in a humidified chamber with 5% CO_2_. All fine chemicals were purchased from Sigma Aldrich and Life Technologies unless otherwise specified.

### Genomic DNA; RNA isolation, Reverse Transcription and Real Time quantitative PCR

Genomic DNA was extracted from K562/ Hek293T cells and used as a template for PCR reactions. The sequence for the PHKB intron 14 was obtained from the ENSEMBLE genome browser. Various primer pairs specific for Hmrhl, Actin, Phkb were obtained from Sigma and used for PCR, cloning, sequencing etc. (Table S1). Total RNA was isolated from cultured cells with the help of TRIzol reagent (Sigma, USA), following manufacturer’s instructions. ~2-3µg of total RNA was reverse transcribed with the help of oligo(dT)_17_ primer and Super Script III/ Revertaid Reverse Transcriptase. 1/20^th^ of the reverse transcription product was used for PCR reaction using gene specific primers. For real-time quantitative PCR (qPCR), the cDNA was diluted to 1:10, added to Sybr green mix (Bio-Rad) and gene specific primers and the reaction was carried out and analyzed in a Biorad Real Time Detection machine.

### Northern Hybridization

The sequence in the central, unique region of Hmrhl from K562 genomic DNA was cloned in pGEM 3Z(+) vector. It was linearized with HinDIII enzyme and antisense digoxigenin-labeled RNA probe were generated using DIG Northern Starter Kit (Roche) according to the manufacturer’s instructions. 2µg of total RNA was loaded on a 2% agarose gel containing 1% formaldehyde and run for 2-3hrs in MOPS buffer. Hybond-N+ membrane (Amersham Bioscience) was used to transfer the RNA on to the membrane through capillary transfer methods overnight. Hybridization was carried out with DIG labeled probe at 60 ˚C overnight. Next day detection was carried out and the membrane was developed according to the DIG Northern Starter Kit instructions (Roche).

### Chromatin immunoprecipitation (ChIP)

K562 cells (10^6^) were fixed in 1% formaldehyde (sigma) for 10 min at RT followed by quenching with 125 mM glycine for 5 min to stop the cross-linking reaction. After washing with PBS containing Protease Inhibitor Cocktail (Roche) twice, cells were spun down and washed with buffer A (20mM HEPES-KOH, pH7.5, 10mM EDTA pH 8.0, 0.5mMEGTA, 0.25% TritonX-100), buffer B (50mM HEPES-KOH, pH7.5, 150mM NaCl, 1mMEDTA pH 8.0, 0.5mMEGTA) for 5-10 min at 4°C and resuspended in buffer C (20mM HEPES-KOH, pH7.5, 1mM EDTA pH 8.0, 0.5mMEGTA, 0.1% SDS and protease inhibitor Cocktail) and allow to sit on ice for 10 min. Samples were sonicated at high intensity with 40 sec on/off cycles in a Bioruptor sonicator (Diagenode) to get fragments in the range of 400-600bp. De-crosslinking was done by adding ProtK and RNase. Samples were diluted 1:5 in IP dilution buffer (20mM Tris pH7.4, 10mM NaCl, 3mM MgCl_2_, 1mM CaCl_2_, 4% NP-40, 1mM PMSF and protease inhibitor). Samples were precleared by binding with 50 µl of protein A beads. An aliquot (50 µl) of soluble chromatin was kept as input. Samples were immunoprecipitated either with the required antibodies (Ab4729 and Ab8895, Abcam) or the pre immune serum as control, O/N at 4°C. Next day 50 µl of protein A beads (Invitrogen) were added to the immune-complex and incubated at 4°C for 2 hrs. Beads were washed 2-3 times sequentially with ChIP wash buffer I (20mM Tris-cl, pH 7.4, 20mM EDTA, 1% Troton X 100, 150mM NaCl, 1mM PMSF), II (20mM Tris-cl, pH 7.4, 2mM EDTA, 1% Troton X 100, 0.1% SDS, 500mM NaCl, 1mM PMSF), and III (10mM Tris-cl, 1mM EDTA, 0.25mM LiCl, 0.5% NP40, 0.5% Sodium deoxy Cholate), then washed with TE and finally chromatin was eluted in 400 µlof elution buffer (25mM Tris-Cl, 10mM EDTA, 0.5% SDS, incubated at 65^0^C for 30 min), DNA was isolated using Promega kit and subjected to qPCR.

### Transfection and Luciferase assay

Clones for luciferase assay were made from K562 genomic DNA using the sequences in the 5’ end of Hmrhl. The inserts were cloned in pGL4.10 (Promoter Vector), pGL4.23 (Enhancer Vector, Promega) between HindIII and Kpn1 sites, upstream of the luciferase gene. The clones were confirmed by sequencing. Primers used are listed in table S1. K562 cells were transfected with the above mentioned clones using lipofectamine reagent. Cells were grown in RPMI supplemented with 10% FBS till they reached confluence (~10^6^ cells in a T25 flask) at which point they were harvested and re-suspended in 1ml of serum free, antibiotic free medium and distributed equally in a 6 well plate which contained medium with 7% FBS. 1µg/ml DNA (clone) was used along with double the amount (v/v) of lipofectamine reagent as transfection solution, which was replaced after 24hours with complete medium and the cells were grown for another day. Cells were harvested after 48 hrs, lysed in 1X reporter lysis buffer for 30 min on ice and transferred to microfuge tubes. Supernatant was collected to which substrate for luciferase was added and readings were recorded in a luminometer.

### siRNA mediated down-regulation of Hmrhl

Four different siRNAs, mapping to the unique, conserved region of Hmrhl were purchased from Sigma 1. 5’-ccaguuacagcaaguacuu-3’; 5’-aaguacuugcuguaacugg-3’ 2. 5’- cauguugcugcuuugguu-3’; 5’-aagccaaagcagcaacaug-3’ 3. 5’-gugacaaagcguucgguau-3’; 5’auaccgaacgcuuugucac-3’ 4. 5’-cuaauccaauauauaaaua-3’; 3’-uauuuauauauuggauuag-3’. A pool of these siRNAs was used for down regulation experiments. K562 cells/ Hek293T cells were transfected with 100nm siRNA per 1.5 ml of the medium (7% FBS), with lipofectamine 2000 reagent in a 6 well plate as described above. Medium was replaced with complete medium (10% FBS) after 24 hrs. Cells were harvested after 48 hrs and RNA isolated and scored for Hmrhl, PHKB, Actin by qPCR. Scrambled siRNA was used for control. Another pool of siRNAs from Dharmacon (Table 1) was also used to down regulate Hmrhl and the effect was examined in both K562 and GM12878 cell lines.

### Immunostaining

Hek293T cells grown on coverslips were fixed for 20 min in 4% paraformaldehyde in PBS, washed with PBS, permeabilized with 0.1% Triton X-100 for 15 min, and blocked with 1% bovine serum albumin (BSA) for 1h. Cells were incubated with primary antibody (β-catenin antibody, Abcam, 1:100 dilution in 0.1% BSA) at room temperature for 45 min followed by three washes with 0.1% PBST (PBS + 0.1% Tween 20) and incubation with Alexa Flour 488 secondary antibody (1:400 dilution in 0.1% BSA) for 45 min at room temperature, washed thrice with 0.1% PBST, stained with 1 µg/ml DAPI (4,6- diamidino- 2-phenylindole), washed and mounted in Dabco (Sigma). Images were acquired in an LSM 510 Meta confocal microscope (Zeiss) and analysed by image analysis software provided by Carl Zeiss.

### Coding Potential and evolutionary conservation analysis

Coding potential of Hmrhl was evaluated with NCBI-ORF finder and CPAT showing the coding probability of ~ 0.99 due to the presence of 1510 bp-long ORF, partially similar (31.1 % similarity in end-to-end global alignment) to the LINE1 ORF2 sequence. To confirm the plausible coding potential of Hmrhl carrying the sequence similar to LINE1-ORF2, PhyloCSF (Phylogenetic Codon Substitution Frequencies) was run showing and strongly advocating that no approximate coding regions or novel coding regions are present. The evolutionary protein-coding potential as determined by Broad Institute’s PhyloCSF data was visualized in UCSC Genome Browser. Furthermore, the absence of Kozak Sequence near the ORF present on HMrhl was confirmed using the online tool “A program for identifying the initiation codons in cDNA sequences”.

For the phylogenetic tree, regions similar to Hmrhl region across 23 species were identified using Ensembl comparative region analysis and the sequences extracted accordingly for each species from Ensembl were used for Multiple Sequence Alignment (MSA) using Clustal Omega leading to the generation of UPGMA tree data in Newick format which was applied for the construction of circular tree in iTOL (Interactive Tree of Life).

For the histone marks at Hmrhl locus, raw data for Histone marks (H3K27ac, H3K4me1, and H3k4me3) was downloaded from UCSC genome browser. After quality processing, FastQ files were aligned against human hg19 genome assembly using TopHat and aligned BAM files were sorted and further used for visualization in IGV Genome Browser.

## Acknowlegments

We thank Suma B. S. of confocal facility and Anitha G. of the sequencing facility at JNCASR. We thank members of M. R. S. Rao laboratory for suggestions. RF would like to thank Dr. Manjira Ghosh, for introduction to the ENCODE database.

## Author Contribution and disclosure of potential conflict of interest

R.F. and M.R.S.R. designed the experiments, analysed the data and wrote the manuscript. R. F., S. R. C. and D. T. R. carried out the molecular and cell biological and experiments. R. F. analysed the ENCODE data. UB performed the bioinformatic analysis for phylogenetic tree, CPAT, PhyloCSF and histone marks. Authors declare no conflict of interest and no competing financial interests.

## References

1. Dunham I, Kundaje A, Aldred SF, et al. ENCODE Project Consortium. An integrated encyclopedia of DNA elements in the human genome. Nature. 2012; 489:57–74.

2. Mercer TR, Dinger ME, Mattick JS. Long non-coding RNAs: insights into functions. Nature reviews genetics. 2009;10(3):155.

3. Nagano T, Fraser P. No-nonsense functions for long noncoding RNAs. Cell. 2011 15;145(2):178–81.

4. Blackwood EM, Kadonaga JT. Going the distance: a current view of enhancer action. Science. 1998;281(5373):60–3.

5. Heintzman ND, Hon GC, Hawkins RD, Kheradpour P, Stark A, Harp LF, Ye Z, Lee LK, Stuart RK, Ching CW, Ching KA. Histone modifications at human enhancers reflect global cell-type-specific gene expression. Nature. 2009;459(7243):108–12.

6. Thurman RE, Rynes E, Humbert R, Vierstra J, Maurano MT, Haugen E, Sheffield NC, Stergachis AB, Wang H, Vernot B, Garg K. The accessible chromatin landscape of the human genome. Nature. 2012;489(7414):75–82.

7. Sakabe NJ, Savic D, Nobrega MA. Transcriptional enhancers in development and disease. Genome biology. 2012;13(1):238.

8. Lovén J, Hoke HA, Lin CY, Lau A, Orlando DA, Vakoc CR, Bradner JE, Lee TI, Young RA. Selective inhibition of tumor oncogenes by disruption of super-enhancers. Cell. 2013;153(2):320–34.

9. Whyte WA, Orlando DA, Hnisz D, Abraham BJ, Lin CY, Kagey MH, Rahl PB, Lee TI, Young RA. Master transcription factors and mediator establish super-enhancers at key cell identity genes. Cell. 2013;153(2):307–19.

10. Hnisz D, Abraham BJ, Lee TI, Lau A, Saint-André V, Sigova AA, Hoke HA, Young RA. Super-enhancers in the control of cell identity and disease. Cell. 2013;155(4):934–47.

11. Kim TK, Hemberg M, Gray JM, Costa AM, Bear DM, Wu J, Harmin DA, Laptewicz M, Barbara-Haley K, Kuersten S, Markenscoff-Papadimitriou E. Widespread transcription at neuronal activity-regulated enhancers. Nature. 2010;465(7295):182–7.

12. Ørom UA, Derrien T, Beringer M, Gumireddy K, Gardini A, Bussotti G, Lai F, Zytnicki M, Notredame C, Huang Q, Guigo R. Long noncoding RNAs with enhancer-like function in human cells. Cell. 2010;143(1):46–58.

13. Mousavi K, Zare H, Dell’Orso S, Grontved L, Gutierrez-Cruz G, Derfoul A, Hager GL, Sartorelli V. eRNAs promote transcription by establishing chromatin accessibility at defined genomic loci. Molecular cell. 2013;51(5):606–17.

14. Lam MT, Li W, Rosenfeld MG, Glass CK. Enhancer RNAs and regulated transcriptional programs. Trends in biochemical sciences. 2014;39(4):170–82.

15. Xiang JF, Yin QF, Chen T, Zhang Y, Zhang XO, Wu Z, Zhang S, Wang HB, Ge J, Lu X, Yang L. Human colorectal cancer-specific CCAT1-L lncRNA regulates long-range chromatin interactions at the MYC locus. Cell research. 2014;24(5):513–31.

16. Kim T, Cui R, Jeon YJ, Lee JH, Lee JH, Sim H, Park JK, Fadda P, Tili E, Nakanishi H, Huh MI. Long-range interaction and correlation between MYC enhancer and oncogenic long noncoding RNA CARLo-5. Proceedings of the National Academy of Sciences. 2014; 111(11):4173–8.

17. Nishant KT, Ravishankar H, Rao MR. Characterization of a mouse recombination hot spot locus encoding a novel non-protein-coding RNA. Molecular and cellular biology. 2004;24(12):5620–34.

18. Ganesan G, Rao SM. A novel noncoding RNA processed by Drosha is restricted to nucleus in mouse. RNA. 2008;14(7):1399–410.

19. Arun G, Akhade VS, Donakonda S, Rao MR. mrhl RNA, a long noncoding RNA, negatively regulates Wnt signaling through its protein partner Ddx5/p68 in mouse spermatogonial cells. Molecular and cellular biology. 2012;32(15):3140–52.

20. Akhade VS, Dighe SN, Kataruka S, Rao MR. Mechanism of Wnt signaling induced down regulation of mrhl long non-coding RNA in mouse spermatogonial cells. Nucleic acids research. 2016;44(1):387–401.

21. Akhade VS, Arun G, Donakonda S, Satyanarayana Rao MR. Genome wide chromatin occupancy of mrhl RNA and its role in gene regulation in mouse spermatogonial cells. RNA biology. 2014;11(10):1262–79.

22. Kataruka S, Akhade VS, Kayyar B, Rao MR. Mrhl lncRNA mediates meiotic commitment of mouse spermatogonial cells by regulating Sox8 expression. Molecular and Cellular Biology. 2017;MCB-00632.

23. Look AT. Oncogenic transcription factors in the human acute leukemias. Science. 1997;278(5340):1059–64.

24. Darnell JE. Transcription factors as targets for cancer therapy. Nature Reviews Cancer. 2002;2(10):740–9.

25. Garraway LA, Lander ES. Lessons from the cancer genome. Cell. 2013;153(1):17–37.

26. Panne D. The enhanceosome. Current opinion in structural biology. 2008;18(2):236–42.

27. Lelli KM, Slattery M, Mann RS. Disentangling the many layers of eukaryotic transcriptional regulation. Annual review of genetics. 2012;46:43–68.

28. Ong CT, Corces VG. Enhancer function: new insights into the regulation of tissue-specific gene expression. Nature Reviews Genetics. 2011;12(4):283–93.

29. Spitz F, Furlong EE. Transcription factors: from enhancer binding to developmental control. Nature Reviews Genetics. 2012;13(9):613–26.

30. Krivega I, Dean A. Enhancer and promoter interactions—long distance calls. Current opinion in genetics & development. 2012;22(2):79–85.

31. Lee TI, Young RA. Transcriptional regulation and its misregulation in disease. Cell. 2013;152(6):1237–51.

32. Goodman RH, Smolik S. CBP/p300 in cell growth, transformation, and development. Genes & development. 2000;14(13):1553–77.

33. Visel A, Blow MJ, Li Z, Zhang T, Akiyama JA, Holt A, Plajzer-Frick I, Shoukry M, Wright C, Chen F, Afzal V. ChIP-seq accurately predicts tissue-specific activity of enhancers. Nature. 2009;457(7231):854–8.

34. Tuan D, Kong S, Hu K. Transcription of the hypersensitive site HS2 enhancer in erythroid cells. Proceedings of the National Academy of Sciences. 1992;89(23):11219–23.

35. Hah N, Murakami S, Nagari A, Danko CG, Kraus WL. Enhancer transcripts mark active estrogen receptor binding sites. Genome research. 2013 May 1.

36. Darrow EM, Chadwick BP. Boosting transcription by transcription: enhancer-associated transcripts. Chromosome research. 2013;21(6-7):713–24.

37. Lai F, Shiekhattar R. Enhancer RNAs: the new molecules of transcription. Current opinion in genetics & development. 2014;25:38–42.

38. Kim TK, Hemberg M, Gray JM. Enhancer RNAs: a class of long noncoding RNAs synthesized at enhancers. Cold Spring Harbor perspectives in biology. 2015;7(1):a018622.

39. Li W, Notani D, Rosenfeld MG. Enhancers as non-coding RNA transcription units: recent insights and future perspectives. Nature Reviews Genetics. 2016;17(4):207–23.

40. Chen H, Du G, Song X, Li L. Non-coding transcripts from enhancers: new insights into enhancer activity and gene expression regulation. Genomics, proteomics & bioinformatics. 2017 Jun 30;15(3):201–7.

41. Boivin V, Deschamps-Francoeur G, Scott MS. Protein coding genes as hosts for noncoding RNA expression. In Seminars in cell & developmental biology 2017. Academic Press.

42. Fatima R, Akhade VS, Pal D, Rao SM. Long noncoding RNAs in development and cancer: potential biomarkers and therapeutic targets. Molecular and cellular therapies. 2015;3(1):5.

43. Tilgner H, Knowles DG, Johnson R, Davis CA, Chakrabortty S, Djebali S, Curado J, Snyder M, Gingeras TR, Guigó R. Deep sequencing of subcellular RNA fractions shows splicing to be predominantly co-transcriptional in the human genome but inefficient for lncRNAs. Genome research. 2012; 22(9):1616–25.

44. Oesterreich FC, Herzel L, Straube K, Hujer K, Howard J, Neugebauer KM. Splicing of nascent RNA coincides with intron exit from RNA polymerase II. Cell. 2016; 7;165(2):372–81.

45. Wallace EW, Beggs JD. Extremely fast and incredibly close: co-transcriptional splicing in budding yeast. Rna. 2017:rna-060830.

46. Terashima M, Fujita Y, Togashi Y, Sakai K, De Velasco MA, Tomida S, Nishio K. KIAA1199 interacts with glycogen phosphorylase kinase β-subunit (PHKB) to promote glycogen breakdown and cancer cell survival. Oncotarget. 2014;5(16):7040.

47. Hehlmann R, Hochhaus A, Baccarani M. Chronic myeloid leukaemia. The Lancet. 2007;370(9584):342–50.

48. Zhou B, Ho S. S, X. Zhu, X. Zhang, N. Spies, S. Byeon, J. G. Arthur, R. Pattni, N. Ben-Efraim, M. S. Haney, et al., “Comprehensive, integrated and phased whole-genome analysis of the primary encode cell line k562,” bioRxiv, p. 192344, 2018.

49. Maurano MT, Humbert R, Rynes E, Thurman RE, Haugen E, Wang H, Reynolds AP, Sandstrom R, Qu H, Brody J, Shafer A. Systematic localization of common disease-associated variation in regulatory DNA. Science. 2012;337(6099):1190–5.

50. Cavalli M, Pan G, Nord H, Wallerman O, Arzt EW, Berggren O, Elvers I, Eloranta ML, Rönnblom L, Toh KL, Wadelius C. Allele-specific transcription factor binding to common and rare variants associated with disease and gene expression. Human genetics. 2016 1;135(5):485–97.

51. Maeda H, Araki K, Kitamura D, Wang J, Watanabe T. Nuclear factors binding to the human immunoglobulin heavy-chain gene enhancer. Nucleic acids research. 1987; 15(7):2851–69.

52. Banerji J, Rusconi S, Schaffner W. Expression of a β-globin gene is enhanced by remote SV40 DNA sequences. Cell. 1981;27(2):299–308.

53. de Villiers J, Schaffner W. A small segment of polyoma virus DNA enhances the expression of a cloned β-globin gene over a distance of 1400 base pairs. Nucleic acids research. 1981;9(23):6251–64.

54. Levinson B, Khoury G, Woude GV, Gruss P. Activation of SV40 genome by 72-base pair tandem repeats of Moloney sarcoma virus. Nature. 1982;295(5850):568–72.

55. Gillies SD, Morrison SL, Oi VT, Tonegawa S. A tissue-specific transcription enhancer element is located in the major intron of a rearranged immunoglobulin heavy chain gene. Cell. 1983;33(3):717–28.

56. Mosthaf L, Pawlita M, Gruss P. A viral enhancer element specifically active in human haematopoietic cells. Nature. 1985;315(6020):597–600.

57. Bories JC, Loiseau P, d’Auriol L, Gontier C, Bensussan A, Degos L, Sigaux F. Regulation of transcription of the human T cell antigen receptor delta chain gene. A T lineage-specific enhancer element is located in the J delta 3-C delta intron. Journal of Experimental Medicine. 1990;171(1):75–83.

58. Hambor JE, Mennone J, Coon ME, Hanke JH, Kavathas P. Identification and characterization of an Alu-containing, T-cell-specific enhancer located in the last intron of the human CD8 alpha gene. Molecular and Cellular Biology. 1993;13(11):7056–70.

59. Speck NA, Renjifo B, Golemis E, Fredrickson TN, Hartley JW, Hopkins N. Mutation of the core or adjacent LVb elements of the Moloney murine leukemia virus enhancer alters disease specificity. Genes & development. 1990;4(2):233–42.

60. Hallberg B, Schmidt J, Luz A, Pedersen FS, Grundström T. SL3-3 enhancer factor 1 transcriptional activators are required for tumor formation by SL3-3 murine leukemia virus. Journal of virology. 1991;65(8):4177–81.

61. Hsiang YH, Spencer D, Wang S, Speck NA, Raulet DH. The role of viral enhancer" core" motif-related sequences in regulating T cell receptor-gamma and-delta gene expression. The Journal of Immunology. 1993;150(9):3905–16.

62. Zaiman AL, Lewis AF, Crute BE, Speck NA, Lenz J. Transcriptional activity of core binding factor-alpha (AML1) and beta subunits on murine leukemia virus enhancer cores. Journal of virology. 1995;69(5):2898–906.

63. Sangle NA, Perkins SL. Core-binding factor acute myeloid leukemia. Archives of pathology & laboratory medicine. 2011;135(11):1504–9.

64. Lewis AF, Stacy T, Green WR, Taddesse-Heath L, Hartley JW, Speck NA. Core-binding factor influences the disease specificity of Moloney murine leukemia virus. Journal of virology. 1999;73(7):5535–47.

65. Hart SM, Foroni L. Core binding factor genes and human leukemia. Haematologica. 2002;87(12):1307–23.

66. Sørensen, K.D., Quintanilla-Martinez, L., Kunder, S., Schmidt, J. and Pedersen, F.S., 2004. Mutation of all Runx (AML1/core) sites in the enhancer of T-lymphomagenic SL3-3 murine leukemia virus unmasks a significant potential for myeloid leukemia induction and favors enhancer evolution toward induction of other disease patterns. Journal of virology, 78(23),13216–13231.

67. Visel A, Minovitsky S, Dubchak I, Pennacchio LA. VISTA Enhancer Browser—a database of tissue-specific human enhancers. Nucleic acids research. 2006;35(suppl_1):D88–92.

68. Ott, C.J., Blackledge, N.P., Kerschner, J.L., Leir, S.H., Crawford, G.E., Cotton, C.U. and Harris, A., 2009. Intronic enhancers coordinate epithelial-specific looping of the active CFTR locus. Proceedings of the National Academy of Sciences, 106(47),19934–19939.

69. Blattler, A., Berman, B.P., Nicolet, C.M., Witt, H., Yao, L., Farnham, P.J. and Guo, Y., 2014. Global loss of DNA methylation uncovers intronic enhancers in genes showing expression changes. Genome biology, 15(9), p. 469.

70. Park SS, Kim JS, Tessarollo L, Owens JD, Peng L, Han SS, Chung ST, Torrey TA, Cheung WC, Polakiewicz RD, McNeil N. Insertion of c-Myc into Igh induces B-cell and plasma-cell neoplasms in mice. Cancer research. 2005 Feb 15;65(4):1306–15.

71. Chapuy B, McKeown MR, Lin CY, Monti S, Roemer MG, Qi J, Rahl PB, Sun HH, Yeda KT, Doench JG, Reichert E. Discovery and characterization of super-enhancer-associated dependencies in diffuse large B cell lymphoma. Cancer cell. 2013;24(6):777–90.

72. Kron KJ, Lupien M, Bailey SD. Enhancer alterations in cancer: a source for a cell identity crisis. Genome medicine. 2014;6(9):77.

73. Corces-Zimmerman MR, Eaton M, Lopez J, Ke N, Fritz C, Olson E, Majeti R, Loven J. Discovery and Characterization of Super-Enhancer-Associated Dependencies in Acute Myeloid Leukemia. Blood. 2014; 124:3539.

